# Peroxisomal Dysfunction Drives Loss of Dynamic Nutrient Responses to Initiate Metabolic Inflexibility During Aging

**DOI:** 10.1101/2025.02.14.638340

**Authors:** Arpit Sharma, Meeta Mistry, Pallas Yao, Yanshan Liang, Sheng Hui, William B. Mair

## Abstract

Aging results in a loss of metabolic flexibility during fasting, characterized by the inability to acutely switch between metabolic substrates and reduced lipid mobilization. However, the drivers of these effects intracellularly remain unclear. Here, in *Caenorhabditis elegans*, we show that loss of coordinated inter-organelle dynamics causally initiates metabolic inflexibility with age. In young animals, peroxisomes emerge as the priming orchestrators of the fasting response, simultaneously governing lipid droplets (LDs) utilization and mitochondrial bioenergetics. With age, peroxisomal priming is lost, leading to mitochondrial fragmentation and impaired dynamic nutrient responses during fasting. Notably, dietary restriction (DR) exerts a rejuvenating effect on peroxisomal function, thereby preserving mitochondrial integrity and promoting longevity. Our study uncovers the expansive network of organelles enabling lipid mobilization during youth, providing critical context to the poorly understood role of peroxisomes in actively maintaining organelle homeodynamics and metabolic flexibility throughout the aging process.

## INTRODUCTION

Cells at their prime are in constant dynamism and maintaining this dynamic response to fluctuating cellular conditions, i.e ‘homeodynamics’, is the truest form of a youthful and functioning system. One fundamental example of homeodynamics is metabolic flexibility, the ability to alternate between different metabolic substrates acutely and appropriately to meet energetic needs of cells, tissues, and the organism^1–5^. With age, metabolic flexibility declines, specifically the responsiveness to fasting and feeding becomes blunted, leading to dysregulated induction of lipid oxidation in older organisms across the evolutionary spectrum^6,7^. Consequently, there is an over-reliance on carbohydrate as fuel, accompanied by increased fat deposition and lipid toxicity^8,9^. These changes collectively contribute to the emergence of age-associated cellular metabolic dysfunction, culminating in chronic metabolic syndrome^10^. Preserving metabolic flexibility is therefore critical for promoting healthy aging and mitigating the onset of age-related metabolic disorders.

Existing studies on loss of metabolic flexibility with age, primarily in obesity or metabolic disorders, have largely focused on inter-tissue hormonal signals that maintain glucose homeostasis, such as insulin production, secretion, and sensitivity^11,12^. Critically, however, much less is known about causal intracellular mechanisms that regulate metabolic flexibility during aging. Under nutrient limiting conditions, the ability to mobilize intracellular lipids from lipid droplets (LDs) is critical for efficient fatty acid (FA) transport and exchange across cells and tissues and thus is the key contributor to organismal metabolic flexibility. Crucially, the ability to mobilize intracellular lipids from LDs declines with age and synchronizes with age-associated metabolic and architectural defects in the key organelles that orchestrate lipid utilization: mitochondria, peroxisomes, and LDs.^13–15^. Recent work has begun to characterize how lipid mobilization is coordinated between these organelle systems during states of metabolic adaptations^16–18^. However, how aging impacts and dysregulates these organelle adaptations to nutrient availability, and whether this leads to intracellular metabolic inflexibility remains unknown.

Here, we identify cellular mechanisms that regulate the metabolic response to fasting in youth that are lost with age, leading to metabolic inflexibility. Using unbiased transcriptomics in *C. elegans*, we show that older animals lose transcriptional flexibility during acute fasting, marked by a specific failure to induce peroxisomal genes. Central nodes regulating peroxisomal protein import, such as PEX-5 rapidly decline with age, rendering peroxisomes import defective, vestigial, and metabolically incompetent. This loss of peroxisome import causally drives pathological LD enlargement and propagation of lipolysis-resistant LDs that are refractory to fasting. Ultimately this leads to a cascade of organelle failure, impairing mitochondrial bioenergetics and initiating defects in fasting metabolism across multiple organelles. Lastly, pro-longevity interventions that invoke states of metabolic adaptation require functional peroxisomes to preserve organelle network homeostasis for longevity. Together, our results highlight the key role of peroxisomes in propagating metabolic adaptation with age and mechanistically identify organellar processes through which peroxisomes alter LD and mitochondrial function during aging to induce metabolic inflexibility.

## RESULTS

### Aging results in loss of transcriptional flexibility and peroxisomal induction during fasting

We characterized metabolic changes in 4 months (young) and 23 months (aged) mice using indirect calorimetric measurements of respiratory exchange ratio (RER). Aging resulted in a global attenuation of *in-vivo* fatty acid oxidation (FAO) reflected in blunted decreases in RER in aged mice during cyclic fasting periods in the light cycle (Figure S1A). To confirm whether aging impedes intracellular lipid mobilization during fasting, we next subjected day 1, (young) and day 8 (aged) adult wild-type (WT) *C. elegans* to acute fasting and quantified lipid abundance. In young animals, lipid quantification using Nile Red (NR) lipophilic dye revealed a clear utilization of lipid stores upon fasting. Contrastingly, aging resulted in a remarkable loss of cellular lipid mobilization during fasting and induced metabolic inflexibility, marked by abundance of lipolysis refractory lipid stores (Figure 1A, S1B). Together revelaing that a global attenuation of lipid mobilization during fasting is universally associated with aging.

**Fig. 1.**
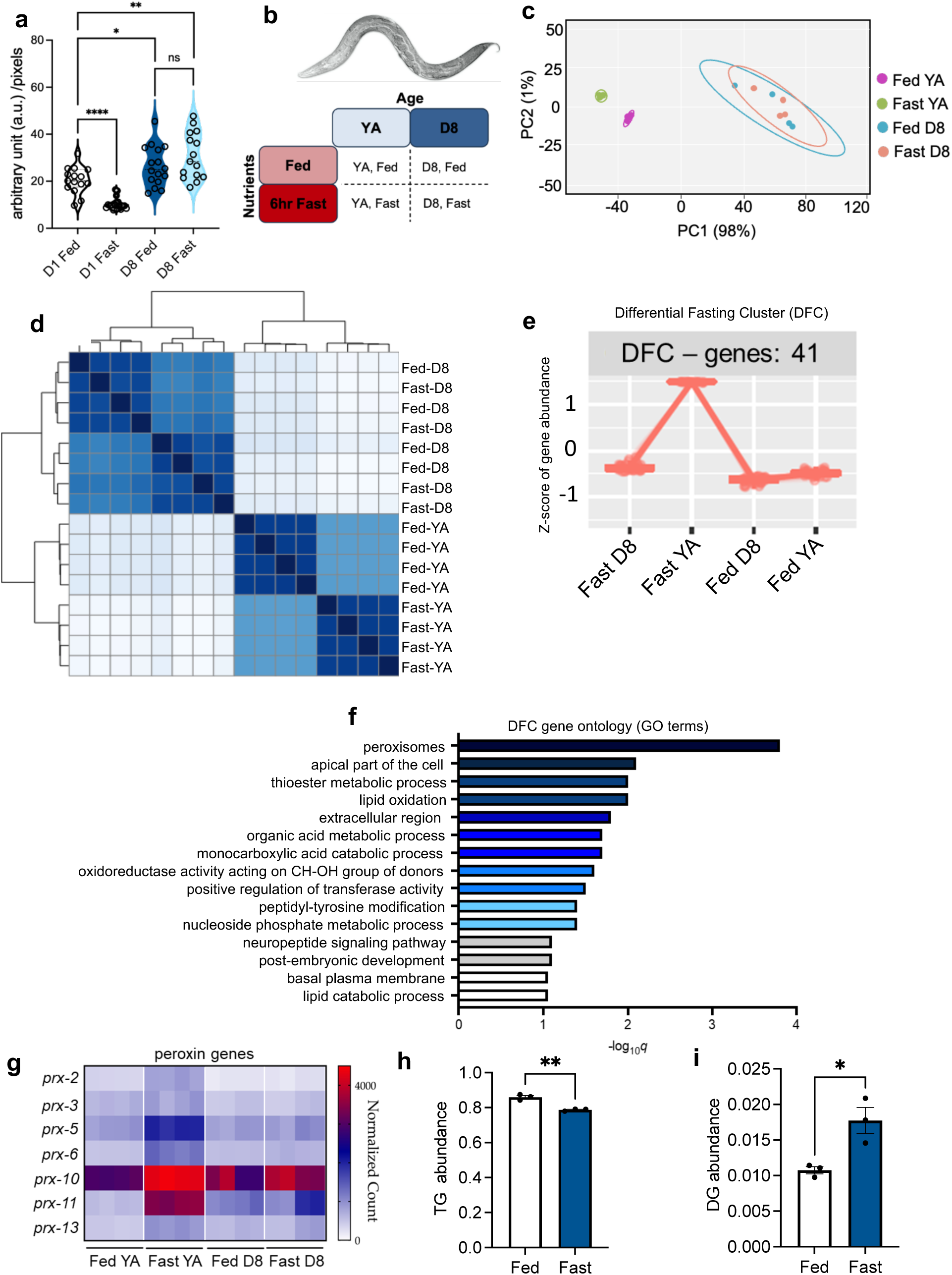
Aging results in organism-wide loss of transcriptional flexibility and peroxisome gene induction. **a**, Quantification of fat stores using Nile Red staining of fixed WT, day 1 or day 8, fed and fasted animals showing induction of metabolic inflexibility with age. (N=3 biological replicates, n=15-18 animals per condition, one way ANOVA). **b,** Experimental design to evaluate changes in fasting responses in young (YA, young adult) and aged (D8, day 8 adult) animals. **c,** Principal component analysis (PCA) of log_2_-transformed and scaled gene expression data for each condition (N=4 biological replicates). **d,** inter-correlation matrix of log_2_-transformed and scaled gene expression data for each condition. **e,** Gene cluster analysis to identify highly specific genes induced with fasting in young but not aged animals, i.e differential fasting cluster (DFC). Differentially expressed genes (DEGs) were first identified using the likelihood ratio test (LRT) and significant genes were obtained using a false discovery ratio (FDR) threshold of 0.01. **f,** Overrepresentation analysis of the gene ontology (GO) biological process terms for genes in the DFC cluster. **g,** heat map representation of pairwise changes in normalized counts of functionally conserved peroxisomal genes (peroxins) in *C. elegans* showing fasting-induction in young but not aged animals. **h,** Triglyceride (TG) abundance in untargeted lipidomic of whole day 1 adult fed and fasted worms using liquid chromatography/mass spectrometry (LC/MS) (N=3 biological replicates, two tailed student’s t test) and normalized to total ion count (TIC). **i,** Diglyceride (DG) abundance in untargeted lipidomic of whole day 1 adult fed and fasted worms using LC/MS and normalized to TIC (N=3 biological replicates, two tailed student’s t test). Data are presented as mean ± SEM (*p<0.05, **p<0.01, ***p<0.001).

To identify cellular mechanisms underpinning this age-associated metabolic flexibility loss, we performed RNA-seq in young (young adult, YA) and aged (day 8 adults, D8) WT *C. elegans* under ad-libitum (AL) fed and 6 hr fasted conditions (Figure 1B). We reasoned that an acute intervention would allow us to identify primary mechanisms induced early on to initiate the physiological fasting response. Principal component analysis (PCA) segregated samples according to age (component 1) and feeding state (component 2) (Figure 1C). In accordance with previous data, a 6 hr fast induced targeted changes in the *C. elegans* transcriptome, while aging induced broad transcriptomic changes, as resolved by the two components^19,20^. While the transcriptional response to an acute fast was robust and distinct in youth, this component distinction with fasting was ablated in aged animals suggesting both, a perturbation in induction of appropriate fasting-responses with age and a loss of transcriptional flexibility in response to fasting (Figure 1C). Crucially, this loss of transcriptional distinction between fasting and feeding with age was not a result of aged animals no longer feeding, as animals still ingest food at day 8 (Figure S1C-D). Similar, to PCA, an intercorrelation analysis revealed that hierarchical clustering could not distinguish between the fed and fasted transcriptomes, specifically in aged animals (Figure 1D). We next validated that in youth, clustering across nutritional status (component 2) is representative of accurate physiological pathways expected to be induced or repressed in response to fasting (Figure 1C). Gene ontology (GO) enrichment analysis verified that young animals broadly induce fatty acid catabolic processes and suppress a wide array of lipid biosynthetic processes in response to a 6 hr food deprivation, accurately adapting to acute nutrient alterations (Figure S1E) ^21^. Since these transcriptional responses to fasting were repressed in aged animals, we set out to directly identify causal genes that are not induced in response to fasting with age.

Using likelihood-ratio test (LRT) clustering, we focused on differentially expressed genes (DEGs) induced by fasting during youth but maximally repressed with fasting in old animals, reasoning that they may mechanistically underlie loss of metabolic flexibility. We identified a highly specific gene cluster, the differential fasting cluster (DFC), of 41 genes with a false discovery rate (FDR) <0.01, enriched with transcripts induced with fasting only during youth, but not with age (Figure 1E). Gene set enrichment analysis on this DFC gene cluster revealed that induction of peroxisome specific genes in response to fasting is specifically lost with age (Figure 1F). Of note, we identified all critical genes related to peroxisomal protein import and lipid oxidation enriched in this DFC cluster, such as *prx-5*, *ech-8*, *daf-22*, and *dhs-28*, all corresponding to the distinct steps in peroxisomal import and FAO (Table S1A and Figure S1F)^22^. Indeed, direct analysis of normalized counts and log_2_FC between the four biological groups confirmed that additionally several conserved peroxins in *C. elegans* show induction with fasting in youth, that is specifically lost in aged animals (Figure 1G, Figure S1G).

Unlike peroxisomal genes, mitochondrial TCA cycle/glycolysis transcripts were predominantly regulated only with age and mitochondrial β-oxidation transcripts were only regulated with fasting, and neither were enriched in the DFC cluster (Table S1A). Together, these data indicate that the blunted response to acute fasting with age is specifically associated with peroxisomal dysfunction and that acute induction of peroxisomal function might precede metabolic alterations in other organelles, such as the mitochondria and LDs. We confirmed that peroxisomal genes transcriptionally induced with fasting in *C. elegans* are also induced in HepG2 cells upon serum and nutrient withdrawal (Figure1G and Figure S1H) demonstrating a conserved engagement of peroxisome pathways across organisms under acute catabolic stimuli^23^. To directly access lipid catabolism during fasting, we performed a lipidomic analysis between WT fed and 12 hr fasted, day 1 animals and found decreased triglycerides (TGs) and a concomitant increase in diglycerides (DGs), indicating lipolysis of stored TGs into DGs in response to fasting (Figure 1H-I). A significant proportion of TGs utilized with fasting contained fatty acid chain lengths corresponding to long chain fatty acids, C>18 (LCFAs) and very long chain fatty acids, C>20 (VLCFAs), both known to be peroxisomal FAO substrates in *C. elegans* (Figure S1I and Table S1B)^24–26^. Together our data suggest that a key primer of acute fasting in youth is activation of peroxisomal metabolism to metabolize stored TGs.

### Peroxisomes become import refractory with age

Peroxisomes are unique organelles that have a specific requirement to import all functional proteins and enzymes inside their lumen. In *C. elegans,* cargo to be imported is recognized by the sole peroxisomal targeting signal 1 (PTS1) and imported via the peroxin regulating peroxisomal protein import, PRX-5^27–29^. Indeed, this critical import regulator, *prx-5,* was the top differentially expressed peroxisomal gene in the DFC cluster (Table S1C). To investigate whether a gradual failure of peroxisome import occurs during aging, we utilized a strain expressing a single-copy GFP-SKL, a green florescent protein fused to the tripeptide PTS1 (Figure S2A-B). We then optimized our previously developed imaging macro “MitoMAPR” (now PeroxiMAPR), to characterize and quantify changes in peroxisomal count and morphology in the metabolically active *C. elegans* intestine (Figure S2C). While peroxisomes exist in various tissues, such as the epidermis (Figure S2D-G), in *C. elegans,* peroxisome import factors like *prx-5* (Figure S2H) *and prx-10* are primarily expressed in the intestine, where metabolic functions analogous to both liver and fat tissues in mammals such as nutrient anabolism and catabolism actively occur^30^.

In young, day 1 animals, we saw a robust sequestration of the GFP signal inside punctate peroxisomal foci (Figure 2A). This localization, although unchanged, increased in variance from day 1 to day 5, reflecting distinct organismal aging trajectories during early-life with varying numbers of import-competent peroxisomes. However, late life animals, at day 8 and day 10, showed a significant loss of GFP peroxisomal foci count, indicating an age-associated failure in peroxisomal protein import and induction of peroxisomal dysfunction (Figure 2A-B). Recent work has shown starvation-induced changes to peroxisomal morphology, specifically tubulation of peroxisomes and their merging with lysosomes, signifying pexophagy in response to nutrient deprivation^31^. We reasoned that changes to peroxisomal import with age might occur simultaneously with alterations in peroxisomal shape. However, in contrast to peroxisomal count, the GFP-SKL did not show changes in peroxisome morphology, i.e length or area with age (Figure 2B-D). To better visualize peroxisomal morphology changes that coincide with merging of acidic lysosomes, we utilized a red fluorescent protein, RFP-SKL strain (Figure S2I), which, unlike GFP, remains unquenched with changes to pH. Similar to the GFP-SKL probe, we identified punctate RFP foci count in young, day 1 animals (Figure 2E). However, starting at age day 5 we saw increasingly tubulated structures, concurrently increasing cytoplasmic RFP fluorescence, lengthening of peroxisomes, and increasing peroxisomal area potentially due to lysosome merging (Figure 2F-H). However, by day 10 we saw highly cytoplasmic, extra-peroxisomal RFP fluorescence and aggregation into clear, non-peroxisomal structures (Figure 2E, 2I). These data corroborated that peroxisomal protein import failure occurs with age, in tandem with peroxisomal morphology changes (Figure S2A)^31,32^.

**Fig. 2.**
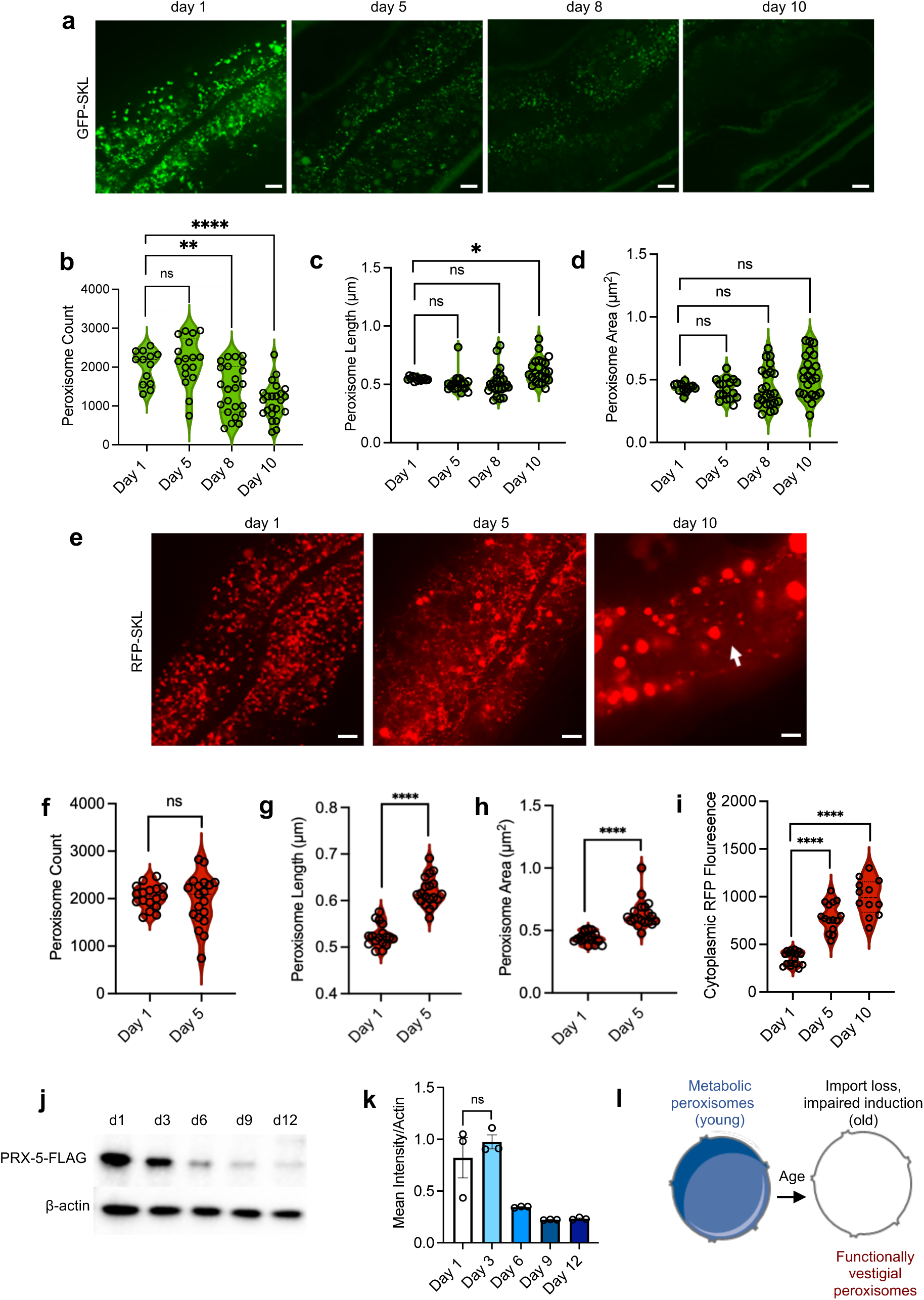
Peroxisomes become functionally impaired during aging. **a**, Fluorescent images of intestinal peroxisome import, identified using GFP-SKL, in the eft3p::GFP-SKL transgenic lines over day 1, 5, 8, and 10 of adulthood. Scale bar = 5 μm. **b-d,** Peroxisome dynamic parameters: count, length (μm), and area (μm^2^) showing a specific decrease in functional peroxisome import of GFP-SKL with age (N=3 biological replicates, n=13-23 animals per condition, one way ANOVA). **e,** Fluorescent images of intestinal peroxisome import, identified using RFP-SKL, in the vha6p::RFP-SKL transgenic lines over day 1, 5, and 10 of adulthood. Scale bar = 5 μm. **f-h,** Peroxisome dynamic parameters: count, length (μm), and area (μm^2^) showing a decrease in functional peroxisome import and alterations in peroxisome length and area with age. RFP fluorescence marks lysosomal targeted peroxisomes (N=3 biological replicates, n=17-19 animals per condition, one way ANOVA). **i,** Cytoplasmic accumulation of RFP-SKL increases with age, quantified from (e). White arrow in day 10 image marks representative area where RFP intensity was quantified, summed, and plotted (N=3 biological replicates, n=12-17 animals per condition, one way ANOVA). **j-k,** Western blot and quantification of endogenously tagged PRX-5::FLAG transgenic lines showing a specific loss of PRX-5 protein levels over day 1, 3, 6, 9, and 12 of adulthood resulting in loss of peroxisome import (N=3 biological replicates, one way ANOVA). **l,** Aging results in a loss of peroxisomal gene induction and peroxisomal import resulting in presence of functionally vestigial peroxisomes. Data are presented as mean ± SEM (*p<0.05, **p<0.01, ***p<0.001).

To investigate whether PRX-5 loss is causal to protein import failure in peroxisomes over age, we generated a separate strain with an N-terminally 3xFLAG-tagged PRX-5. Immunoblotting confirmed a pronounced decline in PRX-5 protein levels from young, day 1 to aged, day 12 animals (Figure 2J-K). Specifically, this robust decline occurred around mid-life, at day 6. Consistent with our transcriptomic and other proteomic observations in *C. elegans*, our data suggest that PRX-5 loss around mid-life contributes to peroxisome import failure with age, resulting in metabolic quiescence characterized by a specific loss of peroxisomal metabolism (Figure 2L)^32^.

### Peroxisome import determines metabolic flexibility

To investigate the metabolic consequences of peroxisome import dysfunction with age, we probed its role in lipid catabolism using mutants with loss of function (*lof*) for import genes *prx-5* and *prx-10*. PRX-10 is known to regulate PRX-5 protein function through monoubiquitylation^33^, and similar to *prx-5* and other peroxins, prx-10 was induced by fasting in our RNA-seq dataset. In WT, *prx-5*, and *prx-10* animals we measured lipid stores under basal, fed conditions using fixed lipophilic NR staining. Supporting a direct role for peroxisome protein import in lipid utilization, fat stores were significantly increased in *prx-5* and *prx-10* mutants (Figure 3A-B). In contrast, loss of *prx-11*, the peroxin regulating peroxisome morphology, but not import, did not affect basal fat stores (Figure S3A-B), highlighting a specific role of peroxisome import in regulating lipid catabolism^34,35^.

**Fig. 3.**
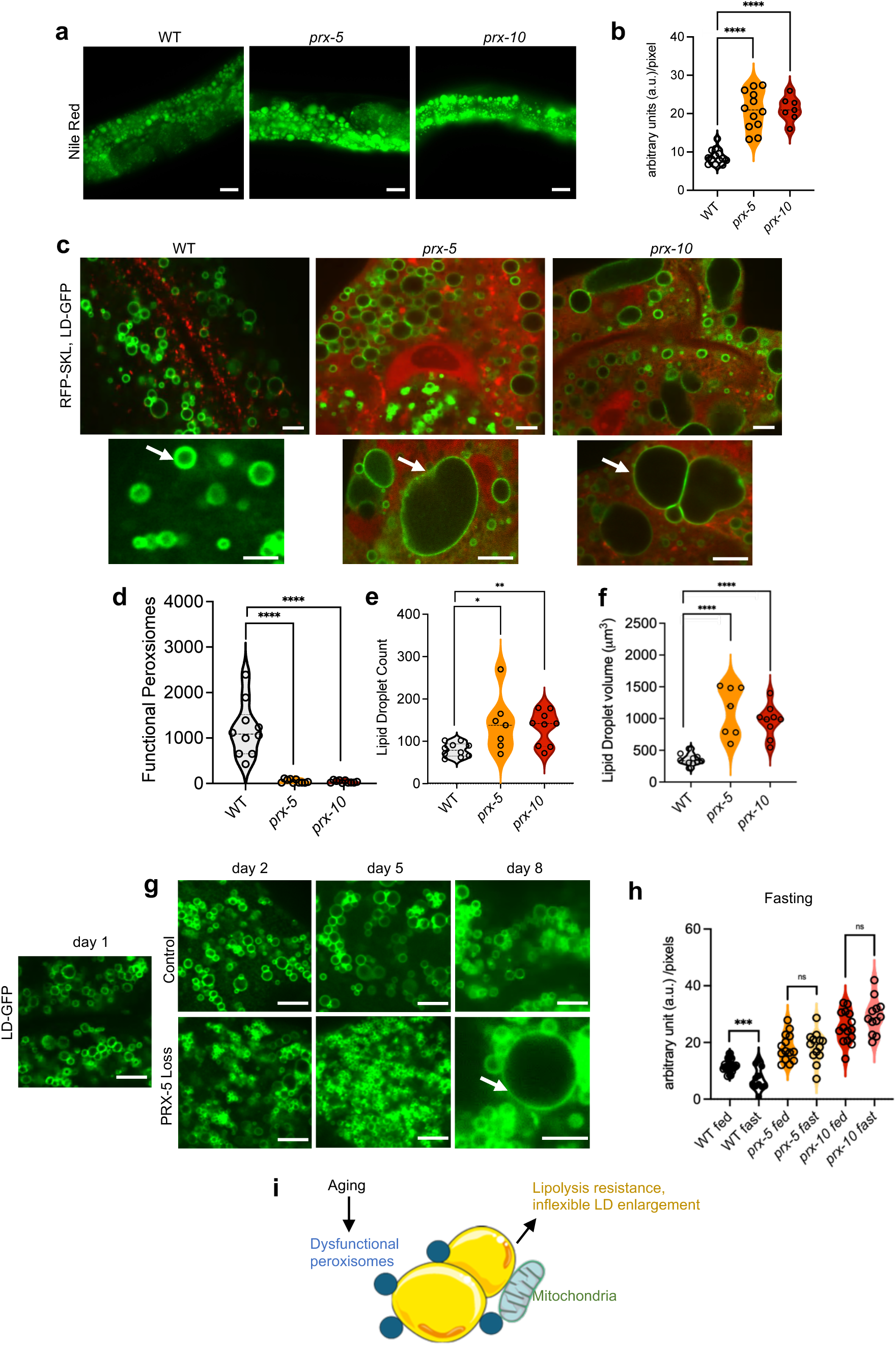
Peroxisome import regulates lipid droplet (LD) dynamics and metabolic flexibility during aging. **a-b**, Labeled imaging and quantification of fat stores using Nile Red staining of fixed WT, *prx-5 (ku517)*, and *prx-10 (ssd68)* animals synchronized at day 1 of adulthood. Scale bar = 20 μm (N=3 biological replicates, n=7-13 animals per condition, one way ANOVA). **c,** Fluorescent images of dual peroxisome-localized RFP and lipid droplet, LD-localized GFP (vha-6p::mRFP-SKL;dhs-3p::dhs-3::GFP) in WT, *prx-5 (ku517)*, and *prx-10 (ssd68)* animals synchronized at day 1 of adulthood. Scale bar = 5μm. Magnified images of LDs are shown underneath each strain. Scale bar = 5μm, arrows highlight LD differences between WT animals and animals lacking peroxisome import. **d,** Quantification of functional peroxisome import of RFP-SKL in either WT animals or animals lacking components of peroxisome import machinery, *prx-5 (ku517)*, and *prx-10 (ssd68)* (N=2 biological replicates, n=7-10 animals per condition, one way ANOVA). **e-f,** Quantification of lipid droplet count and lipid droplet volume (μm^3^) showing increase lipid droplet number and size in *prx-5 (ku517)*, and *prx-10 (ssd68)* animals lacking functional peroxisome import as compared to WT animals (N=2 biological replicates, n=7-10 animals per condition, one way ANOVA). **g,** Fluorescent images of LD-localized GFP (dhs-3p::dhs-3::GFP) imaged at day 1 (baseline) before and after induction of PRX-5 loss over day 2, day 5, and day 8 of adulthood (N=2 biological replicates). White arrow marks grossly oversized LDs only visible over age upon PRX-5 loss specifically. Scale bar = 5μm. **h,** Quantification of fixed Nile Red staining in fed or fasted WT, *prx-5 (ku517)*, and *prx-10 (ssd68)* animals synchronized at day 1 of adulthood showing inflexible LD lipolysis with peroxisome dysfunction. (N=3 biological replicates, n=11-18 animals per condition, two way ANOVA). **i,** peroxisomes regulate LD dynamics through interorganelle lipid metabolism. Data are presented as mean ± SEM (*p<0.05, **p<0.01, ***p<0.001).

Using high-resolution subcellular imaging we next visualized LDs in conjunction with peroxisomes through a strain co-expressing RFP-SKL and DHS-3::GFP, which labels LDs with a GFP fluorophore (Figure S3C). In both *prx-5* and *prx-10* animals, RFP signal was cytoplasmically diffuse, confirming import loss (Figure 3C-D)^16^. Importantly, in accordance with lipid staining, LD abundance was significantly increased in *prx-5* and *prx-10* mutants (Figure 3E). Furthermore, LD volume as a measure of size was significantly increased (Figure 3F), suggesting that increased LD number and their coalescence into supersized structures might result in the presence of LDs with pathological sizes that might become lipolysis resistant. These pathological-sized LDs were absent in *prx-11* null animals confirming that altered peroxisome import, not morphology underlie LD dynamic aberrations and formation of these superstructures (Figure S3D).

To investigate whether peroxisome import loss over age induces pathological LDs, we used targeted protein degradation (TPD) via a degron-tagged PRX-5, initiating acute peroxisome import loss beginning at day 1 of adulthood^36^. While in control animals we identified a uniform increase across LD size, in animals lacking peroxisome import, we identified shifting LD dynamics with age. Around mid-life, day 5, LDs appeared manifold abundant upon PRX-5 loss as compared to control animals. Strikingly around late-life, day 8, we observed emergence of supersized LDs with PRX-5 loss, while these remained absent in control animals (Figure 3G). Next, using lipid staining in conjunction with a feeding-fasting paradigm, we identified that peroxisome import mutants, *prx-5* and *prx-10*, are entirely refractory to fasting-induced lipolysis, displaying the marked characteristic of intracellular metabolic inflexibility i.e., the inability to mobilize lipids (Figure 3H, S3E). Additionally, fasting refractory LDs in the peroxisome import mutants were supersized and irregular (Figure S3E), confirming that peroxisome import is an acute regulator of lipid mobilization with gross effects on LD dynamics.

Notably, as previously observed, although we identified peroxisome tubulation upon fasting (Figure S3F), loss of *prx-11*, which regulates peroxisomal abundance, only affected peroxisome morphology and biogenesis (Figure S3F-I) and did not alter fasting-induced lipolysis (Figure S3E, S3J)^31,37^. Age-associated peroxisome import failure is therefore uniquely linked to the onset of metabolic flexibility and represents a distinct causal regulator of both lipolysis and LD dynamics that is rendered dysfunctional with age (Figure 3I).

### Hallmark mitochondrial defects are induced upon peroxisome import loss

Mitochondria share essential lipid mobilization pathways and biogenesis with peroxisomes^23,38–42^ and the two organelles have defined contact sites^43^. While it has been hypothesized that peroxisomes and mitochondria also age interdependently, the initial step in this inter-organelle failure has remained unclear^42^. We reasoned that peroxisome dysfunction with age might induce metabolic inflexibility by simultaneously promoting mitochondrial failure, resulting in organelle collapse. To dissect the crosstalk between peroxisomes and mitochondria we utilized PRX-5 TPD and monitored mitochondria using a tomm-20(1-49aa)::GFP label (Figure S4A-B). Strikingly, we identified signatures of mitochondrial swelling within 48 hours of PRX-5 degradation, suggesting an acute mitochondrial response upon peroxisomal import loss (Figure 4A). Next, we imaged mitochondria from young, day 1 to aged, day 8 animals with PRX-5 degradation initiated at day 1 of adulthood. As expected, in WT animals we saw age-associated mitochondrial fragmentation, which has been widely observed^15,44–46^. Importantly, in animals lacking peroxisome import, these effects are accelerated. Mitochondrial swelling and blebbing with heterogenous mitochondrial structures appeared as early as day 3, indicating peroxisome collapse accelerates age-related mitochondrial fragmentation (Figure 4B, Figure S4C). Indeed, quantification of mitochondrial count, area, and circularity revealed that animals lacking import-competent peroxisomes had overall fewer mitochondria but increased proportion that were hyper-fragmented and swollen in size (Figure 4C-E). Together these data suggest that defects in mitochondrial biogenesis as well as swelling and fission of existing mitochondria with age are potentially downstream of peroxisome import failure (Figure S4C).

**Fig. 4.**
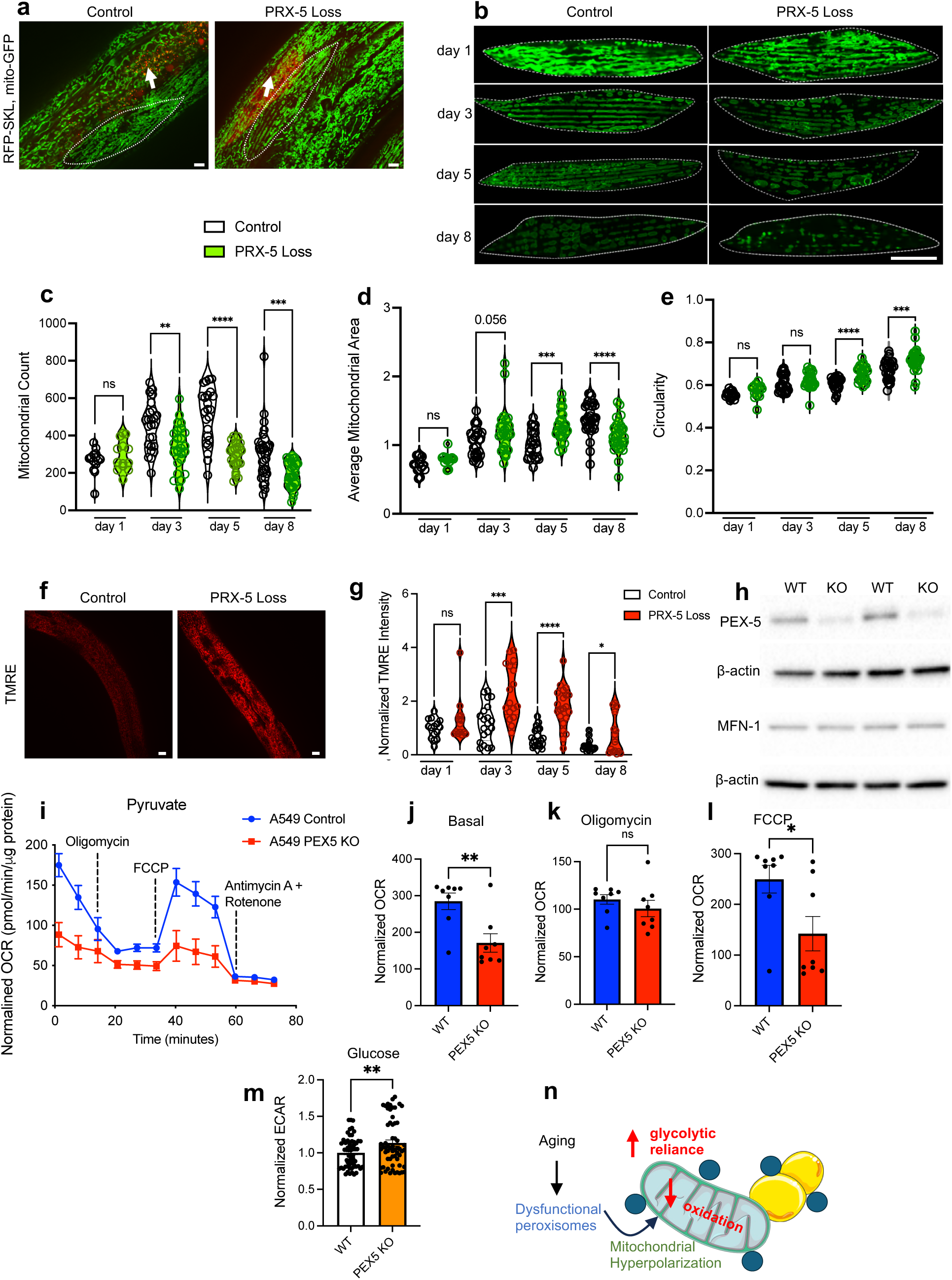
Mitochondrial swelling and biogenetic defects are a consequence of peroxisome import dysfunction during aging. **a**, Representative fluorescent images showing acute mitochondrial dynamic changes upon inducible peroxisomal import loss in transgenic line expressing vha-6p::mRFP-SKL;eft-3p::tomm-20(1-49aa)::GFP to label peroxisomes and mitochondria. Scale bar = 5 µm. (N=3 biological replicates). Arrows mark visible areas where peroxisome import loss phenotype is visible. **b,** Representative fluorescent images showing accelerated swelling of muscle mitochondria with age upon inducible peroxisomal import loss initiated at baseline, day 1 of adulthood and imaged at day 3, 5, and 8 of adulthood subsequently. Scale bar = 5 µm. (N=3 biological replicates). **c-e,** Quantification of mitochondrial dynamic parameters: count, area (μm^2^), and circularity with age in animals with inducible peroxisome import loss initiated at day 1 of adulthood and imaged at day 3, 5, and 8 of adulthood subsequently (N=3 biological replicates, n=13-31 animals per condition, one way ANOVA). **f-g,** Representative fluorescent images and quantification showing mitochondrial hyperpolarization measured by tetramethylrhodamine, ethyl ester (TMRE) retention with age upon inducible peroxisomal import loss initiated at day 1 of adulthood. Images represent day 3 adulthood. Quantification provided over baseline (day 1), and day 3, day 5, and day 8 of adulthood. Scale bar = 10 µm. (N=3 biological replicates, n=14-21 animals per condition, one way ANOVA). **h,** Representative western blot showing PEX5 loss in A549 PEX5 KO cells relative to WT controls. (N=4 biological replicates). **i,** Pyruvate-dependent oxygen consumption rate (OCR) over mitochondrial stress test in WT and PEX5 KO cells show gross mitochondrial oxidative dysfunction in PEX5 KO cells. **j-l,** Quantification of basal, oligomycin inhibited, and FCCP induced maximal respiration over mitochondrial stress test in WT and PEX5 KO cells. (N=3 biological replicates, student’s two-tailed t test). **m,** glucose-dependent extracellular acidification rate (ECAR) in WT and PEX5 KO cells showing increased glycolytic reliance in PEX5 KO cells. (N=2 biological replicates, student’s two-tailed t test). **n,** peroxisomes regulate mitochondrial dynamics and bioenergetics through interorganelle regulation. Data are presented as mean ± SEM (*p<0.05, **p<0.01, ***p<0.001).

To investigate whether peroxisome import failure induces mitochondrial bioenergetic dysfunction and thus mitochondrial swelling, we quantified mitochondrial membrane potential, ΔΨm (MMP) and its attenuation in response to FCCP with tetramethylrhodamine ethyl ester (TMRE) in WT animals (Figure S4D)^47^. While WT animals maintain relatively stable mitochondrial polarity in early adulthood (day 1 and day 3), polarity decreases with age, as previously reported (Figure 4F-G). However, peroxisome import loss induced robust mitochondrial membrane hyperpolarization across age (Figure 4F-G). Critically, ΔΨm varies within a relatively narrow range, −130 to −180 mV and premature deviation from this range, via depolarization or hyperpolarization, induce mitochondrial defects^48,49^. The mitochondrial hyperpolarization observed upon peroxisome import failure with two-fold maximal changes in ΔΨm, represented maladaptive bioenergetics likely to induce age-associated mitochondrial swelling and dysfunction. Importantly, auxin alone did not affect TMRE staining and ΔΨm in WT animals, showing that hyperpolarization of mitochondria is a result of PRX-5 loss and peroxisomal dysfunction (Figure S4E-F).

Since changes to ΔΨm have direct implications on mitochondrial metabolism we next investigated functional alterations induced upon peroxisome import loss. Using WT and PEX5 knockout (KO) epithelial A549 cells we identified a distinct metabolic signature in PEX5-KO cells (Figure 4H, S4G). First, oxygen consumption rate (OCR) on the mitochondrial oxidative source pyruvate was significantly decreased in PEX5-KO cells corroborating perturbed mitochondrial function (Figure 4I). PEX5-KO cells showed a marked reduction in basal and FCCP-induced maximal respiration suggesting significantly reduced oxidative capacity (Figure 4J-L). Second, in contrast to oxidative metabolism, extracellular acidification rate (ECAR) in PEX5-KO cells was significantly greater with glucose as the sole substrate supporting increased reliance on extra-mitochondrial glycolysis and tendency to bypass mitochondrial metabolism for ATP generation in PEX5-KO cells (Figure 4M). Thus, in agreement with our ΔΨm data, PEX5-KO cells harbored bioenergetically perturbed mitochondria with increased reliance on glycolytic metabolism. This metabolic phenotype is also consistent with those observed in cells derived from Zellweger patients with peroxisome biogenesis disorders (PBDs) and rodent models that lack one of the key peroxin genes^50^. Taken together these data indicate that mitochondrial dynamic and metabolic defects commonly associated with age are induced downstream of age-induced peroxisomal import dysfunction (Figure 4N).

### Dietary Restriction (DR) maintains and requires import competent peroxisomes

We have previously shown that a homoeostatic mitochondrial network is critical for DR longevity in *C. elegans*. Therefore we hypothesized that peroxisome import failure through rendering mitochondria fragmented and dysfunctional might make animals refractory to metabolic adaptation required for DR^15^. We first subjected the strain co-expressing RFP-SKL and DHS-3::GFP to DR via bacterial dilution. As expected, animals on DR were significantly longer lived as compared to ad-libitum (AL) fed control animals (Figure 5A). Next, we examined age-related changes to peroxisome protein import in AL and DR treated animals. In young, day 1 adults, we found robust RFP peroxisome import as we previously observed in youth (Figure 5B). However, in aged, day 8 adults, we identified markedly distinct import phenotypes in AL and DR animals. AL fed animals displayed very few import-competent peroxisomes as compared to animals on DR (Figure 5B-C). Surprisingly, aged DR animals displayed cytoplasmic RFP florescence comparable to young AL animals and distinct RFP foci representing import competent peroxisomes (Figure 5B-C). DR therefore maintained import competent peroxisomes. We then asked if DR-longevity might indeed critically require peroxisome function, specifically peroxisome protein import, using PRX-5 TPD. WT animals on DR were significantly longer lived as expected. However, we saw a complete abrogation of DR longevity in animals with induced PRX-5 degradation showing that functional peroxisomes are a central requirement for DR-associated longevity (Figure 5D) and as such why DR maintains increased functional peroxisomes.

**Figure 5.**
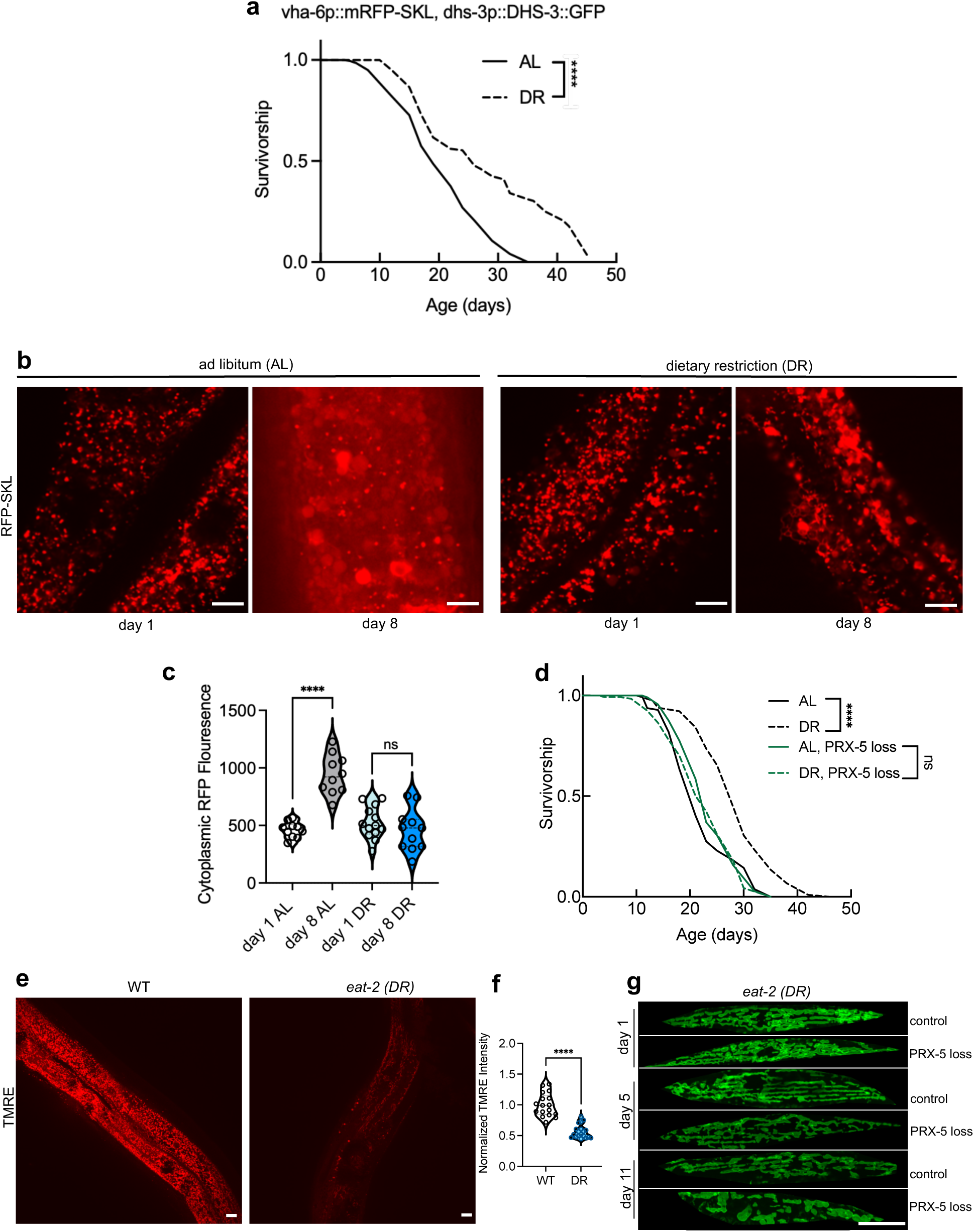
Dietary Restriction (DR) longevity is dependent on functional peroxisomes. **a**, Representative survivorship curve of dual peroxisome-localized RFP and lipid droplet, LD-localized GFP transgenic animals (vha-6p::mRFP-SKL;dhs-3p::dhs-3::GFP) under AL and DR conditions (N=2 biological replicates, n=125 animals per condition, Log rank Mantel-Cox test). **b,** Representative fluorescent images of RFP-SKL in vha-6p::mRFP-SKL;dhs-3p::dhs-3::GFP animals at day 1 and day 8 of AL and DR treatment. Scale bar = 5 µm **c,** Quantification of cytoplasmic RFP fluorescence in vha-6p::mRFP-SKL;dhs-3p::dhs-3::GFP animals at day 1 and day 8 of AL and DR treatment. (N=2 biological replicates, n=10-13 animals per condition, one way ANOVA). **d,** Representative survivorship curve of control animals and animals with PRX-5 degradation under ad-libitum (AL) and DR conditions (N=2 biological replicates, n=125 animals per condition, Log rank Mantel-Cox test). **e-f,** Representative fluorescent images and quantification showing mitochondrial polarization measured by tetramethylrhodamine, ethyl ester is reduced in genetic DR strain *eat-2* as compared to WT animals. Scale bar = 10 µm. (N=2 biological replicates, n=17-19 animals per condition, student’s two tailed t test). **g,** Representative fluorescent images showing mitochondrial fragmentation and swelling of muscle mitochondria with age upon inducible peroxisomal import loss in *eat-2*, DR animals initiated at baseline, day 1 of adulthood and imaged at day 5, and 11 of adulthood subsequently. Scale bar = 5 µm. (N=2 biological replicates). Data are presented as mean ± SEM (*p<0.05, **p<0.01, ***p<0.001).

Lastly, we tested whether the effects of aging and DR on mitochondrial dynamics might depend upon peroxisomal state. We examined ΔΨm in WT and *eat-2 (ad1116)* mutants, a genetic model for DR. In line with previous studies, we found that DR animals have reduced ΔΨm, suggesting mitochondria maybe hypometabolic (Figure 5E-F)^51,52^. ΔΨm hyperpolarization induced upon PRX-5 loss thus may counter a mitochondrial state that is specifically required for DR longevity. To test whether mitochondrial network homeostasis and maintenance of fusion during DR requires functional peroxisomes, we analyzed mitochondrial dynamics in WT and *eat-2* animals at day 1, 5, and 11 of adulthood when adaptations to DR occur (Figure 5G). DR failed to maintain mitochondrial networks when peroxisome function was perturbed; we observed striking mitochondrial fragmentation and swelling in *eat-2* animals upon PRX-5 loss with age supporting the key role of peroxisomes in maintaining a homeostatic mitochondrial network which is amenable to metabolic adaptations for DR longevity. Collectively, our findings suggesting that peroxisome import and function finetune organelle network homeostasis and bioenergetics with age and specifically during DR. Loss of peroxisome function over age consequently is a previously unidentified driver of simultaneous LD and mitochondrial dysfunction and intracellular organelle failure that induces both loss of metabolic flexibility and refractoriness to dietary interventions with age.

## DISCUSSION

Our study identifies mechanisms through which peroxisomes orchestrate organelle network homeodynamics and metabolic flexibility. We find that peroxisomes regulate both dynamics and metabolism of two obligatory organelles involved in cellular lipid mobilization, lipid droplets (LDs) and mitochondria. Temporally, peroxisome function is engaged acutely during fasting to potentially allow for permissive adaptations in both these organelles, but loss of peroxisome function over age renders this organelle network dysfunctional, refractory to metabolic adaptation, and unable to appropriately mobilize lipids. Thus, we identify here, a novel temporal mechanism regulating metabolic flexibility during aging.

Several studies have shown that metabolic flexibility is lost over age, but the precise cellular underpinnings of this loss have remained elusive^2,4,5^. Similarly, the role of peroxisomes in health has remained unclear, with limited studies probing mechanistically how peroxisomal function contributes to cellular and ultimately organismal metabolism. Consequently, existing studies investigating metabolic flexibility have often only identified mitochondria as its unique regulators, driving the cellular response to fasting^53–55^. However, collectively our data show that peroxisomes might play a rheostatic role in regulating both LD and mitochondrial dynamics and function, fine-tuning fasting metabolism, and underpinning organelle dysfunction over age to initiate metabolic inflexibility.

Only recently, work has begun to identify the key role peroxisomes play in regulating lipolysis. PEX5, the mammalian ortholog of PRX5, has been shown to escort the neutral adipose triglyceride lipase (ATGL) onto increased contact points between peroxisomes and LDs to regulate lipolysis^37^. Additionally, adipose-specific loss of peroxisome function has been shown to impair fat thermogenesis, decrease energy expenditure in mice, and increase diet-induced obesity, through specifically modulating mitochondrial fission^56–59^. These recent studies support our model suggesting that peroxisomes might play a previously unrecognized yet critical role in maintaining organismal LD homeostasis through altering mitochondrial form and function.

Here, PRX-5 emerges as a key regulator of metabolic flexibility, specifically enriched in our DFC cluster of transcripts, with other central genes regulating peroxisomal β-oxidation. Indeed, a previous proteomics analysis in *C. elegans* has shown that PRX-5 protein levels decrease with age^32^. Our data suggest a cascade of metabolic alterations emerging from age-associated PRX-5 loss, specifically inducing cellular lipolysis-resistance. PEX5/PRX-5 protein is regulated by several other peroxins, a key one being PEX10. PEX10 monoubiquitinates PEX5 on a conserved N-terminus proximal cysteine residue to enable recycling such that PEX5 can continue import activity^60^. Indeed, cysteine residues have also been shown to become increasingly oxidized with age, rendering them non-functional for regulatory modifications^61^. Therefore, mechanisms both at transcript and protein levels that might alter *prx-5* induction, abundance, and function might contribute to the loss of peroxisome import over age.

A marked phenotype of peroxisome import loss we observed is the increase in LD number and a distinct induction of supersized LDs. LD number has been linked to different effects on health. Previous work has shown that increased LDs are beneficial for longevity, yet LDs have also been shown to accumulate during age and in conditions of systemic metabolic dysregulation^16,62–64^. Together confounding their effects on metabolic health. Our data support that LDs are dynamic organelles interacting with other metabolic counterparts and more than their number, their size, volume, enriched lipids might be indicative of their effect. Thus, the quality of LDs and their dynamic response to changes in nutritional state, rather than their quantity, might be more informative of their role. Supporting this interplay between peroxisomes and LDs, a recent study showed that dietary mono-unsaturated fatty acids (MUFAs) induce beneficial effects on longevity and increase LDs numbers, but specifically require functional peroxisomes^16^. Collectively suggesting that peroxisomal lipolysis might enrich or remove specific lipids that ultimately alter their downstream effect on organismal health, playing a role at par with mitochondrial lipid oxidation.

Indeed, several studies have shown that mitochondria and peroxisomes functionally coordinate lipid oxidation, share common dynamic machinery, and have interlinked biogenesis^41,65,66^. Supporting this intimate functional partnership, we identify that mitochondrial dynamics and bioenergetics are grossly impacted upon loss of peroxisome import over age and importantly animals lacking functional peroxisomes are completely refractory to DR. An underlying regulator of both mitochondrial dynamics and DR-longevity, AMPK, has additionally been shown to consequently increase peroxisome number^15^. Similarly, metformin, which activates AMPK, also increases peroxisome abundance^67^. Together highlighting how conserved signaling cascades known to effect mitochondrial dynamics and metabolism in low-energy states, also might co-regulate peroxisome function. Our data collectively advance existing studies to show not only are import competent peroxisomes maintained during DR, but they are also critical for mitochondrial network homeostasis during DR. Future studies investigating how conserved signaling cascades precisely regulate peroxisome import, dynamics, and metabolism will be key in further dissecting this nuanced interorganelle partnership.

Collectively our findings show that peroxisomes are central orchestrators in the complex interorganelle network regulating lipid homeostasis and that loss of peroxisome function drives age-associated loss of metabolic flexibility. Given the conservation of peroxisome function and lipid metabolism across species, our findings highlight a critical cellular mechanism regulating how efficiently youthful organisms mobilize lipids, originating at the peroxisome. Thus, underscoring the importance of identifying and targeting mechanisms that maintain peroxisome function and plasticity as a novel strategy to restore metabolic flexibility and promote healthy aging.

## Acknowledgments

We thank the Sabri Ülker Center for shared use of their microscopy facility and metabolic chambers. We thank Dr. Suvagata Roy Chowdhury for developing MITOMAPR^68^. We Thank Dr. Anne Lanjuin for critical feedback on the manuscript. AS is funded by NIH/NIA K99AG078343. WBM is funded by NIH/NIA R01AG067106 and R01AG044346.

## Author Contributions

A.S and W.B.M conceived the idea for the study, performed the experiments, analyzed the data, and wrote the manuscript with input from the listed co-authors. M.M at the Harvard Chan Bioinformatics Core helped guide the transcriptomic analysis. P.Y made *C. elegans* strains used to image mitochondria and peroxisomes with GFP fluorophores. Y.L and S.H helped with Lipidomic experiments and data analysis. W.B.M oversaw the project, consulted on its development, and co-wrote the manuscript.

## METHODS

### *C. elegans* strains and husbandry

N2 wild-type, *eat-2 (ad1116),* MH5239 *(prx-5(ku517))*, SOZ259 *(prx-10(ssd68)),* VS10 *(hjIs37 (vha-6p::mRFP-PTS1 + Cbr-unc-119(+)) C. elegans* strains were obtained from the Caenorhabditis Genetic Center, which is funded by NIH Office of Research Infrastructure Programs (P40 OD010440). The strain *vha-7p::mKate2::linker::pxmp-4* was provided by Stéphane G. Rolland. The strain ABR161 (v*ha-6p::mRFP–SKL; dsh-3p::dhs-3::GFP)* was provided by Anne Brunet. Worms were maintained on standard nematode growth media (NGM) seeded with *Escherichia coli* OP50-1 and maintained at 20 °C. The table of strains is available on request.

### Home cage measurements of free-living, whole-body metabolism

C57BL/6 mice were acclimated, young (4 months) and aged (23 months) were acclimated for 24 hrs and then monitered for 60 hrs in an environmentally controlled Sable Systems Promethion Core metabolic cage system fitted with indirect open circuit calorimetry, food consumption momitors, and activity monitors. Respiratory exchange Ratio (RER = VCO_2_/VO_2_) was plotted using CalR Version 1.3 (calrapp.org)

### Microbe strains

OP50-1 bacteria were cultured overnight in Luria-Bertani (LB) broth at 37 °C, after which 100 μl of liquid culture was seeded on nematode growth media (NGM) plates to grow for 2 days at room temperature. Unless otherwise noted, worms were grown at 20 °C on the strain *E. coli* OP50-1 for all experiments. OP50-1 was chosen to remove the need to grow animals in the presence of antibiotics, which can confound organelle functions.

### Solid plate-based dietary restriction assays

Solid dietary restriction assays were performed as previously described^69^. Briefly, ad-libitum (AL) plates were prepared with a bacterial concentration of 100 μl of 10^11^ colony-forming units (CFU) ml^−1^, and dietary restriction (DR) plates with 200 μl of 10^8^ CFU ml^−1^ OP50-1 bacterial concentration. One-hundred microliters of a kanamycin and carbenicillin solution was added after the plates were completely dried (5 mg ml^−1^ and 10 mg ml^−1^, respectively) to arrest bacterial growth at respective AL and DR concentrations. These plates were prepared in advance and stored at 4 °C. 5-Fluoro-2′-deoxyuridine (100 μl of 1 mg ml^−1^ solution in M9) was added on top of the bacterial lawn 24 h before worms were introduced to the plates for lifespans. 5-Fluoro-2′-deoxyuridine was only utilized for lifespans involving dietary restriction and treatment was same across AL and DR conditions. Survival was scored every 1–2 days, and a worm was deemed dead when unresponsive to three taps on the head and tail. Worms were censored due to contamination on the plate, leaving the NGM, eggs hatching inside the adult, or loss of vulval integrity during reproduction. Lifespan analysis was performed using GraphPad Prism.

### *C. elegans* feeding assays

To analyze C. elegans feeding 0.5µm yellow-green YG-GFP beads were used to visualize intake and feeding^70^. Prior to the feeding assay, NGM plates were seeded with 100µl of 0.5µm GFP beads diluted 1:400 in LB broth. Next day, worms were picked from their respective plates and washed three times with 1ml of M9 at 500 rcf and transferred onto unseeded NGM plates. Washed animals from the unseeded NGM plates were then picked onto NGM plates seeded with diluted 0.5µm GFP beads overnight. Animals were allowed to graze on the GFP bead lawn for 3 hours at 20 °C. After 3 hours, animals were rapidly anesthetized using 20 mM tetramisole in M9 and imaged at 20X magnification under 488nm GFP channel to measure feeding and 561nm channel to capture non-specific fluorescence. Mean fluorescence intensity of GFP was then calculated using Image J or FIJI and normalized to animal size to measure direct feed intake into the animals.

### Auxin-inducible total protein degradation (TPD)

Strain PRX-5::degron/AID (auxin-inducible degradation) was generated using CRISPR gene editing and outcrossed. The strain was then crossed into a strain expressing ruby-tagged TIR1 under the control of the somatic eft-3 promoter. To induce degradation of PRX-5 animals were transferred at day 1 of adulthood onto NGM plates containing auxin at a final concentration of 0.15Mm auxin. Control animals were transferred at day 1 of adulthood onto control plates containing an equivalent amount of 100% ethanol. Auxin is light sensitive, and all auxin and control plates were kept away from light to ensure efficacy of auxin treatment.

### Western blots

Around 500 adult worms were used per sample for each appropriate age per replicate. Worms were collected in M9 buffer and washed three times using centrifugation. Liquids were removed after centrifugation, and samples were frozen in liquid nitrogen. For worm lysis, RIPA buffer containing protease inhibitors (Sigma-Aldrich, 8340) and phosphatase inhibitors (Roche, 04906845001) was added to each sample at the same volume as the worm pellet. Worms were lysed via sonication (Qsonica, Q700). Protein concentration was measured using Pierce BCA protein assay kit (Thermo Fisher Scientific, PI23227) following the manufacturer’s instructions. 4× Leammli sample buffer (Bio-Rad, 1610747) was added to denature the proteins, and samples were heated to 95 °C for 5 min. Samples containing 30–40 mg protein were loaded to 10–20% Tris–glycine gels (Thermo Fisher Scientific XP10205BOX) for sodium dodecyl sulfate–polyacrylamide gel electrophoresis. Proteins were transferred to polyvinylidene fluoride membranes (Thermo Fisher Scientific, LC2005) and blocked with 5% non-fat milk powder in TBST. The following primary antibodies were used at the indicated dilutions: monoclonal anti-FLAG M2 antibody produced in mouse (1:1000, Sigma F3165-1MG), β-Actin antibody produced in rabbit (1:1000, Cell Signaling Technology #4967). The secondary antibody was α-mouse IgG HRP-linked (1:1000, Cell Signaling Technology #7076S) and α-rabbit IgG HRP-linked (1:1,000, Cell Signaling 7074). Antibody signals were developed using SuperSignal West Pico PLUS Chemiluminescent Substrate (Thermo Fisher Scientific, 34577), and bands were quantified with Fiji ImageJ.

### Microinjection and CRISPR–Cas9-triggered homologous recombination

All CRISPR edits and insertions required to generate the strains were performed using the previously described CRISPR protocol^71^. Briefly, homology repair (HR) templates were amplified by PCR, using primers that introduced a minimum stretch of 35 bp homology at both ends. Single-stranded oligo donors (ssODN) were also used as repair templates. CRISPR injection mix reagents were added in the following order: 0.375 µl HEPES pH 7.4 (200 mM), 0.25 µl KCl (1 M), 2.5 µl *trans*-activating CRISPR RNA (4 µg µl^−1^), 0.6 µl *dpy-10* CRISPR RNA (2.6 μg μl^−1^), 0.25 μl *dpy-10* ssODN (500 ng μl^−1^) and PCR or ssODN repair template(s) up to 500 ng µl^−1^ final in the mix. Water was added to reach a final volume of 8 µl. Two microliters purified Cas9 (12 μg μl^−1^) added at the end, mixed by pipetting, spun for 2 min at 13,000 rpm and incubated at 37 °C for 10 min. Mixes were microinjected into the germline of day 1 adult hermaphrodite worms using standard methods.

### Fixed Nile Red imaging and analysis

Nile red imaging was performed as previously described^72^. Briefly, at least 100 worms per condition, fed or 12-hour fasted, were buffer washed to remove bacteria and fixed with 40% isopropanol before staining with Nile Red stain for 3 hours at room temperature. Post staining, 3 buffer washes were used to remove excess Nile Red stain. Animals were mounted on a 2% agarose pad and imaged under the bright-field and GFP channel under identical exposure for all conditions imaged. Images were acquired using a Zeiss Axio Imager.M2 microscope equipped with an ApoTome.2 system and an AxioCam MRc camera. Animals were imaged under 10X or 20X zoom to image whole body lipid content and the fluorescence intensity in arbitrary units (a.u.) was quantified using the using the polygon selection tool to select whole animals in ImageJ or FIJI. The flouresecne intensity was normalized for each animal to their selected area and a.u./pixel was obtained. Then, normalized intensity values in a.u./pixels were graphed using GraphPad Prism, and statistical significance was determined using a two-tailed t test. For feeding and fasting Nile Red assays, animals were either ad-libitum fed or fasted overnight to induce lipid utilization.

### Confocal microscopy of mounted worms

For imaging, day 1 adults or appropriate aged adults were mounted on 2% agarose pads and anesthetized with 20 mM tetramisole in M9. For confocal microscopy of organelles such as peroxisomes, mitochondria, and LDs, images were taken in the Sabri Ulker imaging lab using a Yokogawa CSU-X1 spinning confocal disk system (Andor Technology) combined with a Nikon Ti-E inverted microscope (Nikon Instruments). Images were taken using a 100× objective lens, Zyla cMOS (Zyla 4.2 Plus USB3) camera, and 488-nm laser for GFP and 561-nm for RFP.

### Image analysis for peroxisomes and mitochondria

The images were analyzed using macros called PeoxiMAPR for peroxisomes, and MitoMAPR for mitochondria in ImageJ or FIJI^68^. Briefly, representative ROIs were selected from tissue of interest, intestine for peroxisomes and muscle for mitochondria, processed and filtered using the CLAHE plugin with median filter and unsharp mask to increase the local contrast and particle distinctiveness. The ROI was then converted to a binary image to generate a 2D skeleton using the Skeletonize3D plugin. The skeleton image was then dissected using the AnalyzeSkeleton function in FIJI to generate tagged and labelled skeletons. Values obtained from AnalyzeSkeleton functions were used to quantify aspects of peroxisomal and mitochondrial networks respectively. Between images all parameters were kept constant to ensure unbiased analysis through the macros.

### Imaging and analysis of cytoplasmic RFP diffusion

Cytoplasmic localization of RFP-SKL in condition of PRX-5 TPD and aging was quantified from distal intestinal cells. A uniform 4 x 4 µm^2^ area was drawn in distal intestinal cells containing no fluorescence from existing peroxisome punctate and RFP intensity was quantified, summed, and plotted from individual z-sections for each worm to minimize signal masking from from RFP foci (peroxisomes) and to minimize signal bleed through. All analyzed images were obtained using the same exposure time and magnification.

### Imaging and analysis of lipid droplets (LDs)

For LD imaging, the mid-intestinal region was imaged using a 100× objective lens, Zyla cMOS (Zyla 4.2 Plus USB3) camera. The images were taken using the same exposure time/laser power. LD numbers were analyzed in ImageJ or FIJI by applying the same threshold to all images and manually counting the lipid droplets in a 26 × 26 µm^2^ area. The LD diameter was analyzed in ImageJ or FIJI by applying the same threshold to all images and manually measuring the diameter of all the LDs in focus. LD volume estimate measurements were then made from LD diameter.

### Cell culture

HepG2 cells were obtained from American Type Culture Collection (ATCC) and cultured in surface treated dishes in high glucose (4500mg/L) Eagle’s Minimum Essential Medium (EMEM) supplemented with 1Mm glutamine, 10% fetal bovine serum (FBS) and 1% penicillin/streptomycin. For fasting assays, HepG2 cells were cultured in Hank’s Balanced Salt Solution (HBSS) with Ca^2+^ and Mg^2+^ with 10Mm HEPES. Cells were washed twice with PBS at respective fasting collection times, plates were rapidly frozen using liquid nitrogen immersion and RNA was isolated using the Trizol method.

A549 control and PEX5 KO cells were obtained from Christian Metallo’s laboratory at the Salk Institute for Biological Sciences. A549 cells were cultured in ATCC-formulated F-12K medium with 10% FBS and 1% penicillin/streptomycin.

### Quantitative real-time PCR (qPCR)

RNA was isolated from HepG2 cells under fed or respective fasting time points using the Trizol method^53^. cDNA was synthesized from 30 μg of RNA with SuperScript VILO Master Mix (ThermoFisher Scientific, 11755050) following manufacturer’s instructions. 5 ng cDNA was used as template for each qPCR reaction. Each qPCR probe was run along with the control probe as well as no-RT samples (cDNA synthesis reaction set up with RNA but without reverse transcriptase) to be able to detect genomic DNA contamination, if any. No significant genomic DNA contamination was found for any runs. The following probes were used to quantify gene expression on a 96 well plate format using Taqman Universal Master Mix II (Life Technologies, 4440040). Taqman probes used to target each gene of interest are as follows: ACOX1: Hs01074240_m1, PEX5: Hs00165604_m1, PEX6: Hs01122110_m1, PEX10: Hs00538216_m1, PEX12: Hs01558848_g1, PEX13: Hs00968434_m1, PEX14: Hs00992867_m1, PEX19: Hs01005928_m1. Actin was used as the control probe to normalize gene expression: ACTB: Hs99999903_m1.

### High-resolution respirometry assays

A Seahorse Bioscience XF-96 extracellular flux analyzer was used to monitor mitochondrial oxygen consumption similar to how previously described^73,74^. 4 × 10^4^ cells/well were plated onto seahorse cell culture V3 PS plates in their respective culture media overnight and allowed to attach. A three injection mito-stress test protocol in the presence of 10Mm pyruvate in Agilent seahorse media was administered the following day with three replicate measurements taken between each injection. Each replicate consisted of a 1 min mix step, a 1 min equilibration step, and a 3 min measurement step. After basal measurements were acquired, Oligomycin inhibited (1.5µM final concentration, Port A injection), FCCP stimulated (1µM final concentration, Port B injection), and Rotenone+AntimycinA (5µM+4µM final concentration) was used to measure mitochondrial carbohydrate oxidation. Oxygen consumption rate (OCR) was normalized to protein per well quantified using Pierce BCA protein assay kit after respirometry completion.

### Mitochondrial membrane potential ΔΨm (MMP) measurements using TMRE

Tetramethylrhodamine, ethyl ester (TMRE) assays were first performed to validate appropriate MMP staining. A 20Mm TMRE stock in DMSO was used to make a final 10 µM dilution in filtered S-basal buffe. NGM plates prior seeded with 100µl OP50-1 were seeded with 100µl 10 µM TMRE. Appropriate aged animals were transferred to TMRE or control plates overnight. Next day, animals were moved to NGM plates without any TMRE for an hour prior to imaging. Animals were then anesthetized with 20 mM tetramisole in M9 and imaged using a 63× objective lens, Zyla cMOS (Zyla 4.2 Plus USB3) camera, and 561-nm for TMRE fluorescence and 488-nm laser for GFP non-specific background detection. Effect of FCCP on depolarization was also confirmed using the same TMRE staining protocol above but transferring animals onto NGM OP50-1 plates with/without TMRE in addition to 25µM FCCP/control treatment on top of plates. All images were collected at the same exposure time and processed in the same manner. For each image, animal area was selected in ImageJ or FIJI and denoised and deconvolved for better visualization of TMRE stained mitochondria. Fluorescence intensity of TMRE was then quantified and normalized to WT controls to get fold differences in TMRE staining to represent proportional changes in MMP. Normalization did not affect the outcome of the experiments.

### RNA sequencing and differential gene expression analysis

#### RNA collection

Four biological replicates were collected for RNA-seq in the following manner. N2, WT animals were grown on OP50-1 bacteria on 10cm NGM plates to young adult (YA) and day 8 adult (D8) age. On collection, days fed worms were washed from their respective plates with OP50-1 using 15ml M9 + 0.01% Tween. The fed population was washed rapidly three times with M9 buffer using centrifugation. Worms were then reduced to a minimal volume and separated onto NGM OP50-1 seeded (fed) plates and NGM unseeded (fasted) plates and monitored briefly for movement and grazing. After 6 hours of fed and fasted treatment, worms were washed off fed and fasted plates using M9 + 0.01% Tween, washed three times using M9 buffer, pelleted and flash-frozen in 600 μL of Qiazol in liquid nitrogen. Four individual biological replicate samples were thus collected on separate days from separate thaws of N2 strains. Samples were stored at - 80°C.. RNA extractions were performed using Qiagen RNeasy Mini Kit and eluted in RNase free H2O. Samples were sent to the Harvard University Bauer Core Facility for library preparation and RNA sequencing.

#### RNA sequencing

Libraries were prepared using a SciClone G3 NGSx workstation (Perkin Elmer) using the Kapa mRNA HyperPrep kit (Roche Applied Science). Polyadenylated mRNAs were captured using oligo-dT-conjugated magnetic beads (Kapa mRNA HyperPrep kit, Roche Sequencing) from 500 ng of total RNA on a Perkin Elmer SciClone G3 NGSx automated workstation. Poly-adenylated mRNA samples were immediately fragmented to 200- 300bp using heat and magnesium. First strand synthesis was completed using random priming followed by second-strand synthesis and A-tailing. A dUTP was incorporated into the second strand to allow strand-specific sequencing of the library. Libraries were enriched and indexed using 9 cycles of amplification (Kapa mRNA HyperPrep kit, Roche Sequencing) with PCR primers, which included dual 8bp index sequences to allow for multiplexing (IDT for Illumina unique dual 8bp indexes). Excess PCR reagents were removed through magnetic bead-based cleanup using Kapa Pure magnetic beads on a SciClone G3 NGSx workstation (Perkin Elmer). The resulting libraries were assessed using a 4200 TapeStation (Agilent Technologies) and quantified by QPCR (Roche Sequencing). Libraries were pooled and sequenced on one Illumina NovaSeq SP flow cell using paired-end, 75 bp reads.

#### Differential gene expression analysis

Raw reads were examined for quality using FastQC (v0.11.5) to ensure library generation and sequencing were suitable for further analysis. To perform additonal quality checks, all reads were aligned to Ensembl assembly of the C. elegans (PRJNA13758) genome WBcel235_release104 using STAR (v.2.7.0) [57]. Alignments were checked for biases, evenness of coverage, rRNA content, genomic content of alignments, complexity and other quality checks using a combination of FastQC, Qualimap, and MultiQC. To quantify the abundance of reads corresponding to each transcript, alignment and quantification was performed using Salmon (1.4.0). Differentially expressed genes (DEGs) were identified using the Wald test, and significant genes were obtained using a false discovery rate (FDR) threshold of 0.01. Comparisons of the differential gene expression between the various conditions was performed using DESeq2 from tximport with a p-vaule cutoff of 0.01. Principal component analysis was performed using DESeq2.

#### Likelihood Ratio Test (LRT)

To derive gene clustering information to identify subsets of genes that all change with fasting in young but not aged animals LRT cultering was used. Differentially expressed genes obtained using a false discovery rate of >0.01 were separated into clusters based on similar expression profiles across young adult-fed (YA-fed), young adult-fasted (YA-fast), day 8-fed (D8-fed), and day-8 fasted (D8-fast) using a function from the DEGreport (v1.20.2) package. Gene lists for each cluster were used as input to the R bioconducator package clusterProfiler (v3.14.3). As a quality check, normalized counts were also manually plotted for select genes in clusters to ensure changes in fold change followed appropriate clustering profile.

### Lipidomics

Methyl-tert-Butyl Ether (MBTE) extraction was used for fast extraction of total lipid extracts from 2000 fed and 12-hour fasted N2 worms each. A 12 hr fast was chosen to induce changes to the lipidome subseuquent to acute transcriptional changes. Three biological replicates of N2 worms borh fed and fasted were collected for the lipidomics analysis. The extracted lipids were reconstituted in solvent volumes proportional to protein concentrations for normalization. 100ul of supernatant was used for LC-MS based lipidomic profiling. Lipid peaks passing QC were used for LipidSearch based annotation.

### Extended Data Figures

**Fig. 1.**
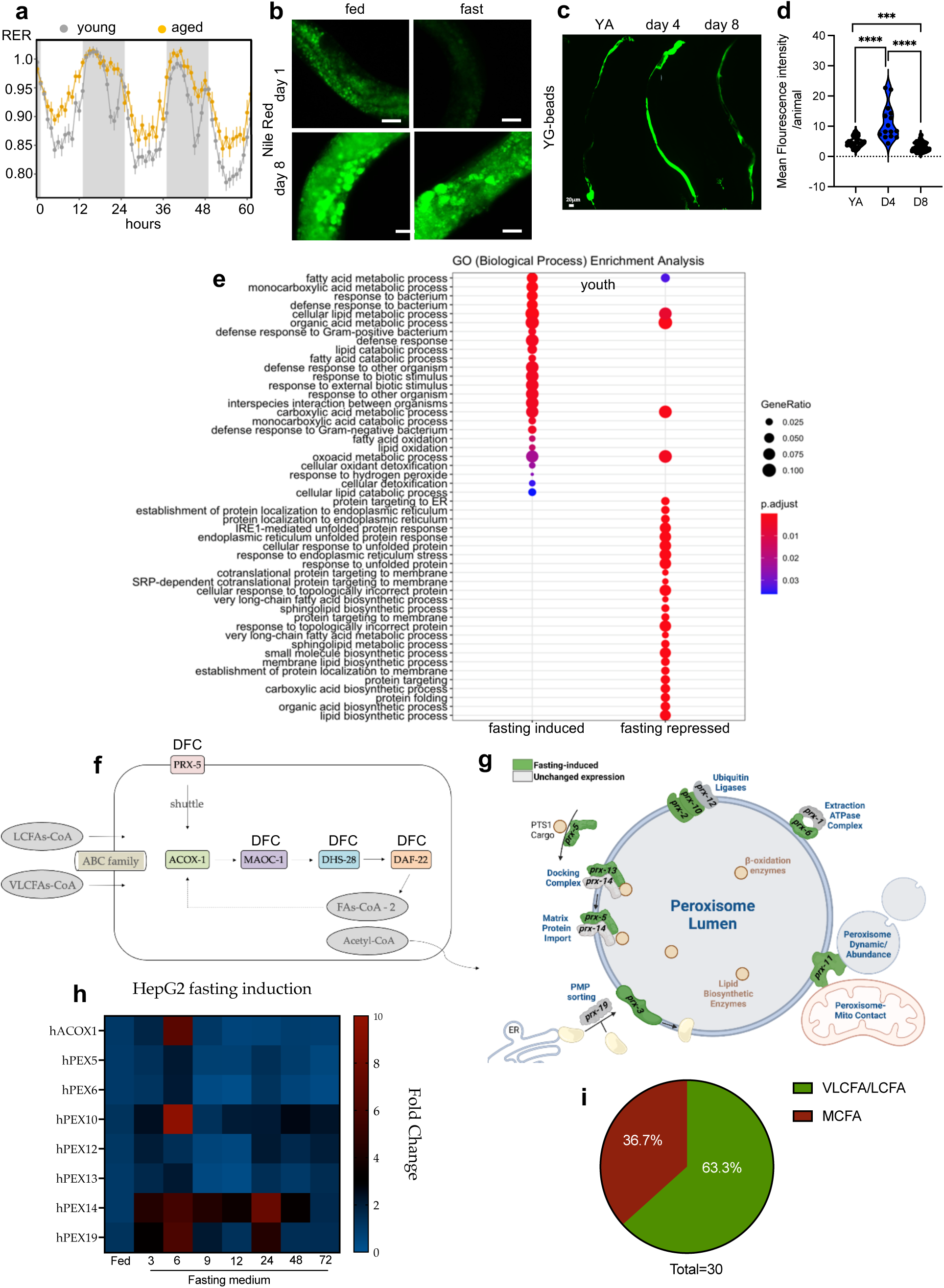
Aging is associated with an attenuation of peroxisome gene induction with fasting. **a, Respiratory Exchange Ratio (RER) of WT young (n=8, age = 4 months) and aged (n=8, age = 23 months) mice. b,** Labeled imaging of fat stores using Nile Red staining of fixed WT, day 1 or day 8, fed and fasted animals showing induction of metabolic inflexibility with age. Scale bar = 20 μm. (N=3 biological replicates, n=15-18 animals per condition, one way ANOVA). **c-d,** Fluorescent images and quantification measuring food intake using yellow green (YG) fluorescent beads in young adult (YA), day 4 adult (D4), and day 8 (D8) adult animals showing aged animals pump and eat. Scale bar = 20 µm. (N=2 biological replicates, n=15-35 animals per condition). **e,** Overrepresentation analysis of the GO biological process terms for genes induced and repressed with fasting. **f,** Schematic of peroxisomal β-oxidation. Genes *prx-5*, *ech-8* (part of multifunctional enzyme along with *maoc-1*), *dhs-28*, and *daf-22* belong to enriched DFC cluster, induced with fasting in young but not aged animals. “DFC” represents genes in the differential fasting cluster **g,** Schematic showing pairwise comparison of fasting-induced peroxin genes (green) and their role in peroxisomal function. Every function comprised a fasting-induced peroxin gene. (N=4 biological replicates). **h,** Heatmap showing fold change in gene expression of conserved peroxins with fasting in HepG2 cells. (N=3 biological replicates). **i,** Fractional clustering of triglycerides (TGs) that decrease with fasting in day 1 adults shows significant peroxisomal processed TGs containing long chain and very long chain fatty acids (LCFAs/VLCFAs) as compared to medium chain fatty acids (MCFAs). (N=3 biological replicates). Data are presented as mean ± SEM (*p<0.05, **p<0.01, ***p<0.001).

**Fig. 2.**
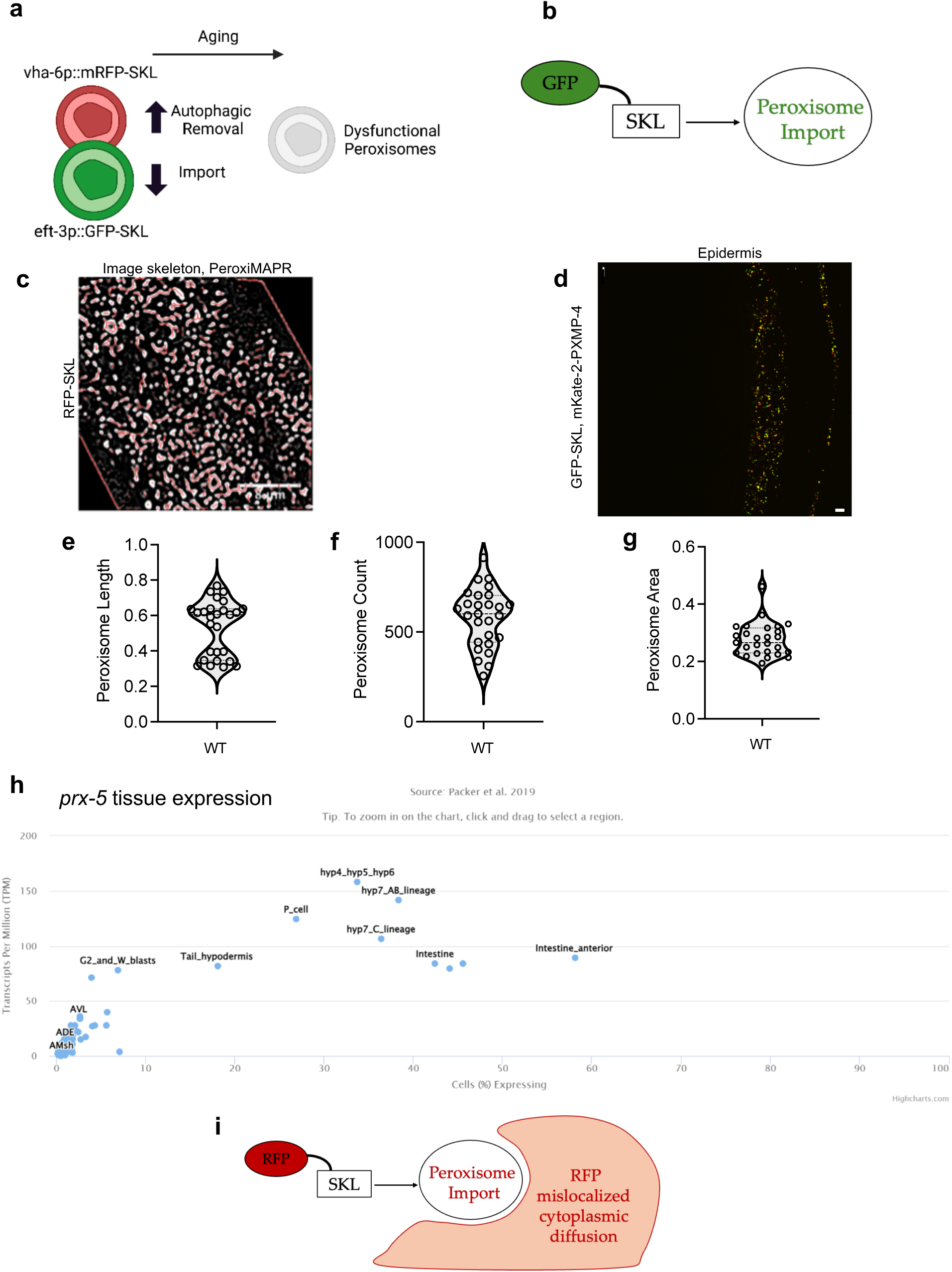
Peroxisomal protein Import is predominant in *C. elegans* intestine. **a-b**, Schematic of different transgenic strains containing GFP-SKL (eft3p::GFP-SKL) and RFP-SKL (vha6p-mRFP-SKL) utilized to measure peroxisome import and autophagic removal with age respectively. Higher tissue penetration and photostability of RFP was used to quantify and evaluate cytoplasmic RFP-SKL diffusion. **c,** Representative skeletonized image of RFP-SKL in the intestine used to quantify import competent peroxisomes using PeroxiMAPR. Scale bar = 8 µm**. d-g,** Fluorescent image and peroxisomal dynamics (count, length, and area) in *C. elegans* epidermis, imaged using GFP-SKL and mkate2-PXMP-4 co-marker transgenic strain to show presence of peroxisomes in several tissues. Scale bar = 5 µm. (N=2 biological replicates, n=27 animals).**h,** Tissue expression atlas of *prx-5* showing predominant expression in the metabolic tissue, the intestine. **i,** Schematic of RFP-SKL (vha6p-mRFP-SKL) utilized to simultaneously visualize peroxisome import, cytoplasmic RFP diffusion, and peroxisomal dynamics.

**Fig. 3.**
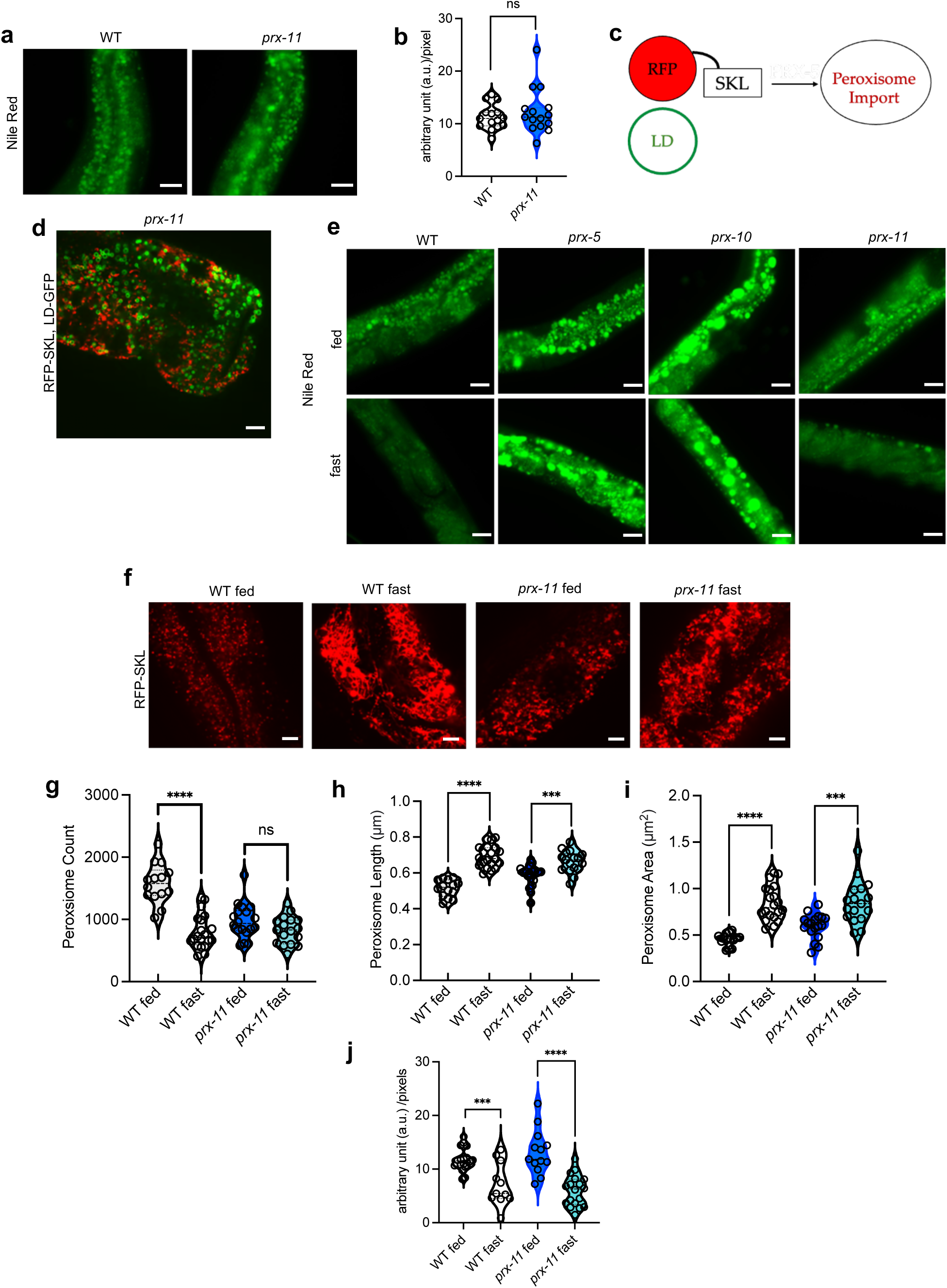
Peroxisome import machinery and not dynamics regulates fasting metabolism. **a-b**, Labeled imaging and quantification of fat stores using Nile Red staining of fixed WT and *prx-11* animals synchronized at day 1 of adulthood. Scale bar = 10μm. (N=3 biological replicates, n=15 animals per condition, student’s two tailed t test). **c,** Schematic of peroxisome-localized RFP and lipid droplet, LD-localized GFP transgenic (vha-6p::mRFP-SKL;dhs-3p::dhs-3::GFP) imaging strain utilized to image peroxisomes and LDs simultaneously. **d,** Representative florescent image of LDs using vha-6p::mRFP-SKL;dhs-3p::dhs-3::GFP strain in the *prx-11* background regularly shaped LDs comparable to WT animals. Scale bar = 5 μm. (N=3 biological replicates). **e,** Representative images of fat stores using Nile Red staining of fixed WT, *prx-5*, *prx-10* and *prx-11* animals fed and fasted animals synchronized at day 1 of adulthood. WT and *prx-11* animals mobilize lipids upon lipolysis, whereas *prx-5* and *prx-10* animals do not. Scale bar = 20 μm. **f-i,** Dynamic changes in peroxisomal morphology with fasting in WT day 1 adult animals are blunted in *prx-11* mutants. Peroxisomal biogenesis is reduced in *prx-11* mutants and fasting-induced increase in length and are reduced. Scale bar = 5 μm. (N=3 biological replicates, n=14-20 animals per condition, two way ANOVA). **j,** Quantification of fixed Nile Red staining in fed and fasted WT and prx-11 animals shows both strains perform fasting-induced lipolysis. (N=3 biological replicates, n=11-18 animals per condition, two way ANOVA). Data are presented as mean ± SEM (*p<0.05, **p<0.01, ***p<0.001).

**Fig. 4.**
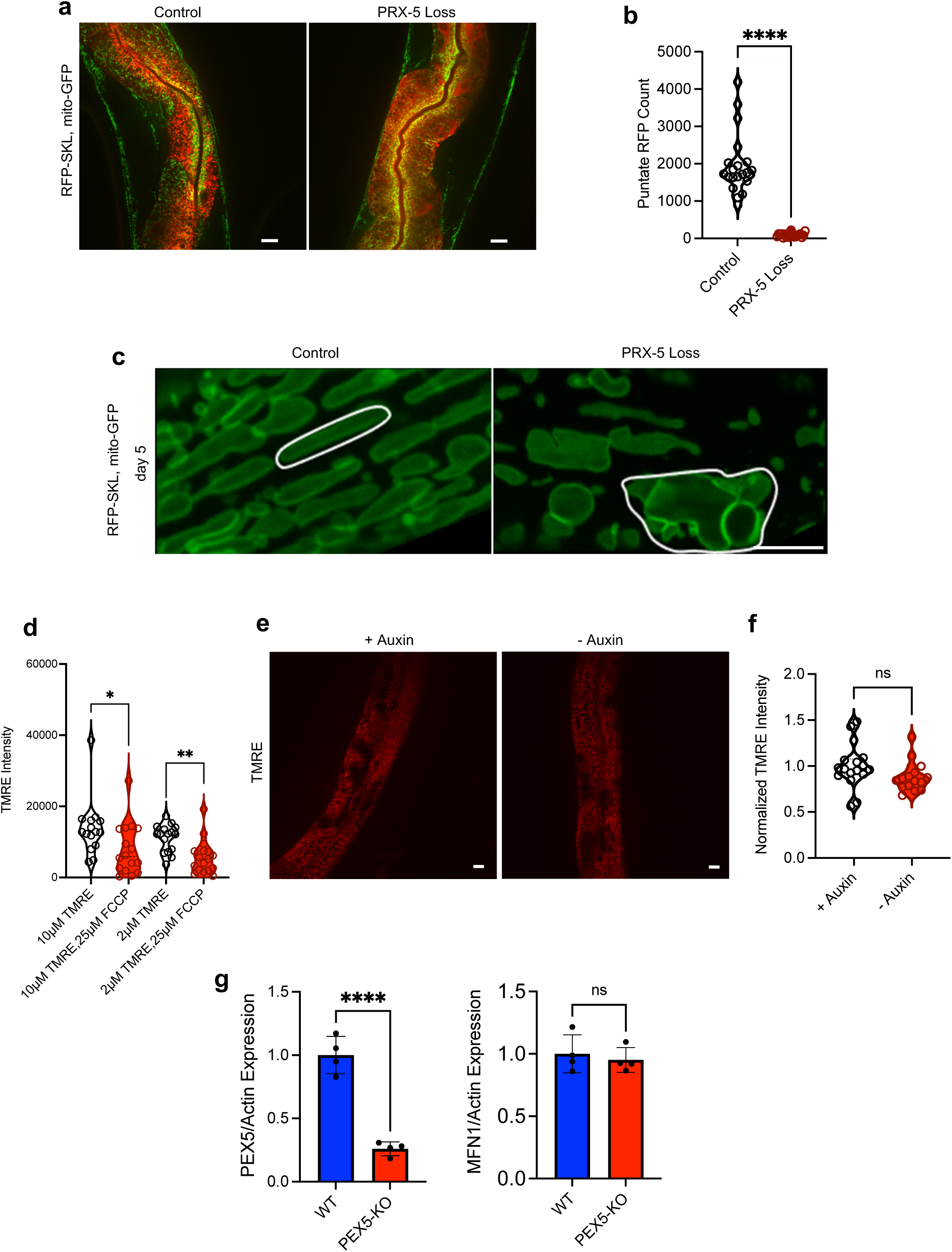
Peroxisomal import loss induces mitochondrial defects during aging. **a**, Fluorescent images showing the effect of PRX-5 loss on peroxisome import. RFP puncta become cytoplasmic upon induced PRX-5 loss. Scale bar = 10µm. **b,** Quantification of punctate peroxisomes upon inducible loss of PRX-5 protein. (N=3 biological replicates, n=20 animals per condition, student’s two tailed t test) **c,** Magnified images of GFP labelled mitochondria in day 5 control and PRX-5 degraded adults showing gross alterations and mitochondrial swelling upon PRX-5 loss. White encircled mitochondria highlight common defects. Scale bar = 5 µm **d,** Quantification of by tetramethylrhodamine, ethyl ester (TMRE) intensity in two TMRE concentrations (10μM and 2μM) with and without 25μM FCCP treatment to establish that TMRE readily stains polarized mitochondria, and its intensity decreases with FCCP treatment. (N=2 biological replicates, n=15-17 animals per condition, student’s two tailed t test). **e-f,** Representative fluorescent images and quantification showing auxin treatment alone does not affect mitochondrial polarization in WT animals. Scale bar = 10 µm (N=2 biological replicates, n=15-17 animals per condition, student’s two tailed t test). **g,** Quantification of PEX5/Actin and MFN1/Actin expression in WT and PEX5-KO A549 cells. (N=4 biological replicates, student’s two tailed t test). (*p<0.05, **p<0.01, ***p<0.001).

